# H3K9me2 genome-wide distribution in the holocentric insect *Spodoptera frugiperda* (Lepidoptera: Noctuidae)

**DOI:** 10.1101/2021.07.07.451438

**Authors:** Sandra Nhim, Sylvie Gimenez, Rima Nait-Saidi, Dany Severac, Kiwoong Nam, Emmanuelle d’Alençon, Nicolas Nègre

## Abstract

**Background:** Eukaryotic genomes are packaged by Histone proteins in a structure called chromatin. There are different chromatin types. Euchromatin is typically associated with decondensed, transcriptionally active regions and heterochromatin to more condensed regions of the chromosomes. Methylation of Lysine 9 of Histone H3 (H3K9me) is a conserved biochemical marker of heterochromatin. In many organisms, heterochromatin is usually localized at telomeric as well as pericentromeric regions but can also be found at interstitial chromosomal loci. This distribution may vary in different species depending on their general chromosomal organization. Holocentric species such as *Spodoptera frugiperda* (Lepidoptera: Noctuidae) possess dispersed centromeres instead of a monocentric one and thus no observable pericentromeric compartment. To identify the localization of heterochromatin in such species we performed ChIP-Seq experiments and analyzed the distribution of the heterochromatin marker H3K9me2 in the Sf9 cell line and whole 4^th^ instar larvae (L4) in relation to RNA-Seq data.

**Results:** In both samples we measured an enrichment of H3K9me2 at the (sub) telomeres, rDNA loci, and satellite DNA sequences, which could represent dispersed centromeric regions. We also observed that density of H3K9me2 is positively correlated with transposable elements and protein-coding genes. But contrary to most model organisms, H3K9me2 density is not correlated with transcriptional repression.

**Conclusion:** This is the first genome-wide ChIP-Seq analysis conducted in *S. frugiperda* for H3K9me2. Compared to model organisms, this mark is found in expected chromosomal compartments such as rDNA and telomeres. However, it is also localized at numerous dispersed regions, instead of the well described large pericentromeric domains, indicating that H3K9me2 might not represent a classical heterochromatin marker in Lepidoptera.

## Introduction

The nuclear organization of the genome into chromatin is a hallmark of eukaryotes. DNA is wrapped around histone proteins to form nucleosomes that constitute basic units of chromatin (Luger et al. 1997). Two types of chromatin are classically described based on the compaction of nucleosomes along the genome. The euchromatin represents “open” and less compacted chromatin structures and is usually associated with active gene transcription. On the other hand, heterochromatin designates regions of the chromosomes that are more compact, with a higher nucleosome density (Heitz, 1928). Genes within heterochromatin are regarded to be transcriptionally repressed.

Several types of heterochromatin have been described. Constitutive heterochromatin (c-Het), contrary to facultative heterochromatin, remains persistently compacted despite cell cycle and developmental stages, environmental states or even studied species (Dillon 2004; Allshire et Madhani 2018; Janssen, Colmenares, et Karpen 2018). It is usually located at important chromosomal features such as telomeres, rDNA loci and pericentromeric regions (Riddle et al. 2011). Those regions are usually gene poor and transcriptionally silenced (Dillon 2004; Allshire et Madhani 2018). Understood as a genomic safety guard from transposons (Janssen, Colmenares, et Karpen 2018), c-Het associated regions are often enriched in repeated sequences such as satellite DNA and transposable elements. The transcriptional silencing by c-Het is due to its compaction which is achieved by chemical modifications of histones. The classical marker of c-Het in all eukaryotes is the post translational methylation of the lysine 9 of Histone H3 (H3K9me) (Kharchenko et al. 2011; Liu et al. 2011; Ho et al. 2014). This methylation mark deposited by SET domain proteins such as Su(var)3-9 (Rea et al. 2000) and G9a (M. Tachibana et al. 2001; Makoto Tachibana et al. 2002; Kondo et al. 2008) is recognized by chromo-domain containing proteins belonging to HP1s family (Minc et al. 1999; Lachner et al. 2001; Maison et Almouzni 2004; Lomberk, Wallrath, et Urrutia 2006). HP1 proteins assemble as homodimers to form the ultrastructural 3D compaction detected by cytology (Verschure et al. 2005). This compaction impairs the binding of other DNA associated proteins such as transcription factors and RNA polymerases, which explains the repressive effect of heterochromatin. These properties of c-Het have been well described in model organisms, from yeast to mammals and thus are thought to be conserved.

With developments of sequencing, biochemical methods and growing interest for non-model organisms (Tagu, Colbourne, et Nègre 2014), the classical c-Het features are being reconsidered (Grewal et Jia 2007). Underlying DNA sequences can show rapid turnover, a fact particularly true for centromeres (Henikoff, Ahmad, et Malik 2001). Depending on cell cycle phases, nascent non coding RNAs from telomeres and pericentromeres contribute to the regulation of their biology (Bierhoff, Postepska-Igielska, et Grummt 2014; Allshire et Madhani 2018). H3K9me distribution has been shown to vary and dynamic apposition of histone marks has also been reported outside of primary c-Het regions (Wen et al. 2009) in heterochromatin “islands” (Riddle et al. 2011; Lee et al. 2020). In human and mouse, interstitial domains called LOCKS, spanning several Mb, dynamically switch to heterochromatin state, marked by Lysine 9 methylation, in specialized cells, supposedly to limit pluripotency (Wen et al. 2009; Madani Tonekaboni, Haibe-Kains, et Lupien 2021). Beside these controlled variations, c-Het can unpredictably change in terms of associated proteins or even DNA sequences. This causes several defects in development or represent molecular basis of hybrid incompatibilities between close species (Ferree et Barbash 2009; Hughes et Hawley 2009; Johnson 2010; Crespi et Nosil 2013; Satyaki et al. 2014).

While most studies on c-Het have been performed on monocentric species, a particular case of c-Het dynamic is found in holocentric organisms. Their chromosomes have no cytological hypercompacted regions and possess dispersed centromeres instead of unique ones per chromosome (Schrader 1935). Classical heterochromatin compartmentation and properties are thought to be absent. Holocentrism has been described in several plants, in some nematodes and some insect orders (Wolf 1996; Drinnenberg et al. 2014). Phylogenetic analysis indicates that holocentric species might have derived several times from monocentric species by convergent evolution (Melters et al. 2012; Escudero, Márquez-Corro & Hipp 2016). This is supported by different centromeric molecular signatures between holocentric species. In monocentric species, major 150bp satellite forming centromeres are packaged in CenH3 rich nucleosomes that are encompassed by peripheral H3K9me2/3 regions (Melters et al. 2013; Drinnenberg et al. 2014). But, except for the plant *Rhynchospora pubera* (Marques et al. 2015), described holocentric species have lost those centromeric repeated sequences. CenH3 is present in nematodes (Gassmann et al. 2012; Steiner et Henikoff 2014) and in plants (Zedek et Bureš 2016; Oliveira et al. 2020) but has been lost in other holocentric insects (Drinnenberg et al. 2014). A recent study proposed that in Lepidopteran, CenH3 function has been replaced by H3K27me3, a facultative heterochromatin mark (Senaratne et al. 2021). More importantly H3K9me2 signal surrounding centromeres is thought to be lost in these organisms, unlike holocentric *C.elegans* (Steiner et Henikoff 2014). Previous studies conducted on Lepidopteran showed nonetheless that H3K9me2 is still associated to repeated DNA at rDNA loci and sexual chromosomes (Stanojcic et al. 2011; Borsatti et Mandrioli 2013).

In this paper, we aim to clarify H3K9me2 heterochromatin distribution in *Spodoptera frugiperda* (Lepidoptera: Noctuidae), a crop pest causing severe damages to plants at larval stage. Since the *S. frugjperda* distribution area has recently been extended from the American continent to a worldwide invasion (Goergen et al. 2016), there is an urge to understand its adaptive potential when confronted with new ecosystems. In particular, chromatin properties could influence phenotypic plasticity in response to environmental conditions (Simola et al. 2016; Gibert et al. 2016). *S.frugiperda* constitutes also an emergent epigenetic model organism with published reference genomes for different strains and cell lines (Kakumani et al. 2014; Nandakumar, Ma, et Khan 2017; Gouin et al. 2017; Nam et al. 2020; L. Zhang et al. 2020), histone modifications and non-coding RNAs being previously characterized (Stanojcic et al. 2011; d’Alençon et al. 2011; Moné et al. 2018) as well as a growing body of RNA-Seq data (Orsucci et al. 2020). Another advantage lies in the well-established Sf9 cell line, derived from *S. frugiperda* ovarian tissues, providing non limiting material for biochemical assays (Vaughn et al. 1977). We performed H3K9me2 ChIP-Seq and RNA-Seq on two different samples: Sf9 cell lines and whole 4th instar larvae (L4). In both samples, we confirmed the association of H3K9me2 at c-Het domains such as telomeres, rDNA locus and satellite sequences that might represent vestigial centromeres. We found a strong association of H3K9me2 with repeat elements as well as gene bodies. And we show that H3K9me2 enrichment at these elements is not associated with transcriptional repression or activation, raising the question of its role in chromosomal organization in holocentric lepidopteran.

## Material and Methods

### *Spodoptera frugiperda* rearing and Sf9 cell line maintenance

L4 insects have been raised in controlled laboratory conditions of 16h:8h light/dark photoperiod cycle, ~40 % mean hygrometry and ~24°c temperature. The insects derived from pupae individuals collected in 2001 in Guadeloupe. This laboratory population corresponds to previously published reference genome assemblies (Gouin et al. 2017; Nam et al. 2020). Immortalized Sf9 cell line derived from *S. frugiperda* ovarian tissues (Vaughn et al. 1977). The cell line was acquired from ATCC (https://www.atcc.org/products/crl-1711) and has been cultured following the manufacturer protocol recommendations.

### Western blot

Chromatin extracts were prepared from Sf9 cells and L4 insects as described below fro the ChIP procedure. Total proteins have been first quantified by colorimetric Bradford method and equivalent quantities were used for a 15% SDS/PAGE. After migration, proteins from the gel were transferred onto a PVDF membrane. The membrane was incubated overnight with mouse monoclonal H3K9me2 antibody (Abcam 1220) and revealed with an ECL kit.

### Immunofluorescence on Sf9 cell lines

Sf9 cells were grown to confluence with standard Schneider medium, then scraped and collected in 50 mL Falcon tubes. Cells were then diluted up to 3.105 cells/ml and 1 mL was used for each immunostaining condition for 1 well of a 12-well plate. Plates with round glass coverslips and 1 mL of cell dilution were then placed at 28 °C for 4 h, allowing the cells to sediment on the coverslip. The culture medium was then removed, plates washed with 1X PBS and fixed with paraformaldehyde 4%, 20 min at room temperature. Coverslips were rinsed twice with PBS before processing with immunostaining.

For immunostaining, coverslips in the plates were permeabilized with 1 mL of PBS 1X + Triton 1% per well for 30 min at room temperature. The solution was removed, and the cells were blocked with PBS 1X + BSA 1% for 30 min at room temperature. The solution was removed and staining with the primary antibody (anti-H3K9m2, Abcam 1220) diluted in PBS-BSA 0.1 % was performed. We used 1/100 and 1/200 dilutions (50 μL well) and we kept a control non-treated well. Incubation was done 1 h at room temperature. Primary antibody was rinsed twice in PBS 1X before adding the secondary antibody (anti-mouse Alexa568) diluted 1/500 in PBS 1X – BSA 0.1% (~100 μL per well) 30-45 min at room temperature in the dark. Still in the dark, coverslips were rinsed once in PBS 1X, then incubated with DAPI (1mg/mL diluted 1/1000 per condition) for 5 minutes and rinsed again. Coverslips were then mounted on a microscopy slide with 1 drop of ProLong Antifade Mountant (ThermoFischer Scientific) and after 30min, sealed with transparent nail polish. Slides were kept in the dark at 4°C until observation with an Apotome microscope (Zeiss).

### ChIP-Seq and RNA-Seq

We performed ChIP-Seq following an adapted protocol by Nègre et al. 2006 on 4th instar whole larvae and Sf9 cell culture.

Briefly, biological samples are being crushed in a douncer, in presence of 1% formaldehyde and incubated for 15 minutes to allow protein-DNA crosslinking. Crosslinking was quenched by the addition of 225mM glycin. Chromatin was then fragmented using a Bioruptor sonicator (Diagenode). An aliquot of sonicated chromatin at this stage is used as the Input sample.

For the immunoprecipitation, sonicated chromatin is incubated for 4 h with the primary antibody at 4°C, then with Protein-A sepharose beads (CL4B) for again 4 h. Beads were then centrifuged and washed. Chromatin was eluted from the beads with a 50mM Tris-HCl buffer with 1%SDS and 10mM EDTA. To reverse the cross-linking, this immuno-precipitate was incubated overnight at 65°C and then 2-3h at 50°C with proteinase K. DNA was purified from this precipitate with 500μL of phenol-chloroform and 55μL of 4M LiCl. The DNA from the aqueous phase is precipitated with 100% ethanol and the pellet dried and resuspended in water. Chromatin was prepared from 50 L4 larvae pool and 50 mL of confluent cells per IP condition. Two biological replicates for input and ChIP conditions have been systematically produced except for larvae (1 replicate only). Genetic material sequenced in 50bp single-end (Illumina-seq technology).

We also performed RNA-Seq experiments from Sf9 cell lines. RNA was purified using TRIzol (Invitrogen) following manufacturer’s instructions and sent to Genewiz for strand-oriented, mRNA sequencing. Previously published RNA-Seq data for L4 (Orsucci et al. 2020) genomes were used to perform combined analysis.

Datasets are available in ArrayExpress (https://www.ebi.ac.uk/arrayexpress/) with the following accession numbers: E-MTAB-6540 for L4 RNA-Seq; E-MATB-10686 for Sf9 RNA-Seq and E-MTAB-10721 for ChIP-Seq experiments.

### H3K9me2 ChIP-Seq analysis

Raw reads from fastq files have been filtered using cutadapt software and trimmed for remaining adapters, low quality (phredscore < 35) and short length sequences (<40nt). Corresponding samples alignements were performed using stringent parameters with Bowtie2 (--endtoend --verysensitive, Langmead et al. 2009) against Sf9 (Nandakumar, Ma, et Khan 2017) and L4 genomes (Nam et al. 2020). Samtools have been used for sample concatenation (Li et al. 2009) after Deeptools -bigwigcorrelate Pearson analysis (Ramírez et al. 2016). H3K9me2 whole genome peak detection has been assessed with MACS2 callpeak using the following parameters: --broadpeak -f BAM -g 340000000 -n --down-sample (Y. Zhang et al. 2008). Peaks were annotated using Bedtools software (--intersect) after functional annotation of genomic elements. Nucleotide peak comparison between the two models have been performed using online D-genies software (Cabanettes et Klopp 2018).

H3K9me2 plot and heatmap over genomic location of interests were produced from Deeptools (computeMatrix and plotHeatmap option).

Integrative Genomic Viewer (IGV) was used for annotation and sequencing track visualization (Thorvaldsdóttir, Robinson, et Mesirov 2013).

### Genome functional element annotation

We used published *S. frugiperda* transcriptome (Legeai et al. 2014) for Sf9 and L4 gene annotations using Scipio software (Keller et al. 2008). It predicts gene exons by detecting intron/exon junctions with the BLAT algorithm. CDS were reconstructed using Bedtools -- groupby function. Introns were annotated by subtracting CDS-exons positions (Bedtools -- subtract).

RNA-Seq data were filtered, concatenated and aligned on the genome with the same methods as described for H3K9me2 ChIP-Seq samples. Expression analysis has been performed with Stringtie software (Pertea et al. 2015; parameter: stringtie ‘bam files’ -G genereference.gtf -e). Output in transcript per millions (TPM) was converted to log2 to determine active pool genes (log2(TPM)>1) from inactive ones (log2(TPM)<=1). Since UTRs detections failed after several software tests (such as Maker, exUTR, getUTR or KLEAT), we arbitrarily defined 3’UTRs to be 500pb long succeeding last exon (bedtools flank -l 0 -r 500 -s), 5’UTRs to be 200 bp long preceding first exon (bedtools flank -l 200 -r 0 -s) and promoters to be 1000bp long preceding 5’UTRs.

Satellite DNA has been detected using TandemRepeatFinder (Benson 1999), Repeatexplorer (Novák et al. 2013) and Repeatmasker on both genomes. Major 150bp satellite regions were shuffled using Bedtools --shuffle option.

We characterized repeat DNA with consensus sequence inferior or equal to 10 base pairs as microsatellites. Those comprising [10-100]bp where qualified as minisatellites. Finally, consensus sequence satellite of over 100 bp were annotated as satDNA.

We used blastn with default parameters to annotate transposable elements in Sf9 and L4 genomes, using previously determined transposable elements consensus sequences from *S. frugiperda* (Gouin et al., 2017) annotated by the REPET pipeline (Quesneville et al., 2005, sequences with less than 70 % homology were filtered out), rDNA copies (with a minimum of 1000 nucleotides length (Gouin et al., 2017)) and telomeres (Gong et al. 2015) in *Spodoptera* genus.

### Statistic and graphic productions

Barplot, scatterplot, histogram and Venn Diagram were plotted using R software. Student t-test and Chi-square test were performed using R software. Gene ontology enrichment has been performed using BLAST2GO annotations followed by Fisher exact test analysis (Conesa et al. 2005).

## Results

### 1. H3K9me2 genome-wide distribution in Sf9 and L4 genomes

To investigate the presumptive genome-wide localization of heterochromatin and its dynamic in *S.frugiperda* we performed ChIP-Seq experiments against the H3K9me2 chromatin mark in Sf9 cells as well as in L4 caterpillars. We first tested the specificity of the mouse anti-H3K9me2 antibody (ab:1220) by immunofluorescence experiments in Sf9 cells. We observed a strong nuclear signal that disappears during metaphase (**Supplementary Fig. 1**). This is a well-known characteristic of this mark, which is usually shielded by Histone H3 Serine 10 Phosphorylation during mitosis (Duan et al. 2008; Jeong et al. 2010; Poleshko et al. 2019). We also performed western-blot on chromatin samples showing a 17kDa band corresponding to the mark in both samples (**Supplementary Fig. 2**). We then performed ChIP-Seq experiments following an adapted cross-linking procedure for our insect model (see **Methods**). For each experimental condition (input vs. ChIP-Seq, Sf9 vs. L4) we sequenced two biological replicates, except for L4 input.

After removing short and bad quality sequences, reads were mapped against respective reference genomes for Sf9 cell lines (Nandakumar, Ma, et Khan 2017) and L4 *S.frugiperda* larvae (Nam et al. 2020) (**Table 1**). The alignment rates range between 86.44% and 96.53%. Interestingly, both Sf9 and L4 ChIP-Seq shows a higher level of multimappers (reads mapping more than once) compared to input (**Table 1**). Indeed, a strong association between H3K9me2 and repeated sequences has been found previously described in Lepidopteran (Stanojcic et al. 2011; Borsatti et Mandrioli 2013). Multimappers number is higher in Sf9 than in L4. This can be explained by Sf9 polyploidy (Jarman-Smith et al. 2002) or independent Sf9 and L4 sequencing and bioinformatic genome assemblies (Nandakumar, Ma, et Khan 2017; Nam et al. 2020).

**Table 1:**
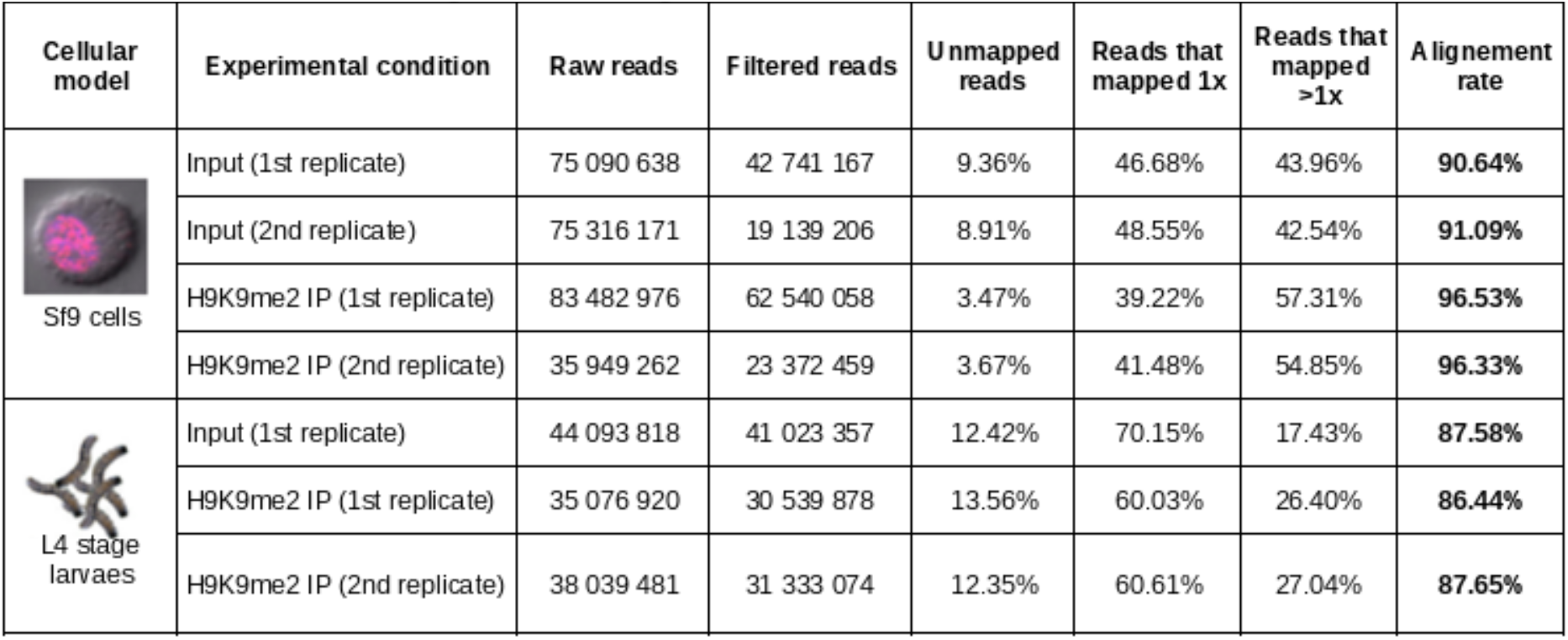
Input and H3K9me2 ChlP-seq sequencing properties in Sf9 cells and L4 larvaes. Raw, filtered reads and their alignements usin Bowtie2 software.

| Cellular model | Experimental condition | Raw reads | Filtered reads | Unmapped reads | Reads that mapped 1x | Reads that mapped >1x | Alignment rate |
| --- | --- | --- | --- | --- | --- | --- | --- |
| Sf9 cells | Input (1st replicate) | 75 090 638 | 42 741 167 | 9.36% | 46.68% | 43.96% | **90.64%** |
|  | Input (2nd replicate) | 75 316 171 | 19 139 206 | 8.91% | 48.55% | 42.54% | **91.09%** |
|  | H9K9me2 IP (1st replicate) | 83 482 976 | 62 540 058 | 3.47% | 39.22% | 57.31% | **96.53%** |
|  | H9K9me2 IP (2nd replicate) | 35 949 262 | 23 372 459 | 3.67% | 41.48% | 54.85% | **96.33%** |
| L4 stage larvae | Input (1st replicate) | 44 093 818 | 41 023 357 | 12.42% | 70.15% | 17.43% | **87.58%** |
|  | H9K9me2 IP (1st replicate) | 35 076 920 | 30 539 878 | 13.56% | 60.03% | 26.40% | **86.44%** |
|  | H9K9me2 IP (2nd replicate) | 38 039 481 | 31 333 074 | 12.35% | 60.61% | 27.04% | **87.65%** |

When we tested the similarity of reads enrichment per genomic positions between samples (**Supplementary Fig. 3** and detailed in **Methods**) we observed that Pearson correlation R coefficient is higher between ChIP samples (R=0.65 for Sf9; R=0.85 for larvae) than between Input samples (R=0.44 in larvae), a result consistent with independently sonicated samples. Since ChIP-Seq and input data segregate separately in both samples (**Supplementary Fig. 3**), we concatenated corresponding replicates and analyzed H3K9me2 enrichment by comparing ChIP-Seq over input. Respectively 35 596 and 30 382 peaks of enrichment were found in Sf9 and L4 samples. Half of them were small peaks comprising 0 to ~1000bp, whereas the other half formed larger genomic domains (comprising between ~1000bp to several 10 000 bp, **Fig. 1A-B**). Interestingly, H3K9me2 covers 13.8±0.02% and 12.6±0.03% of total Sf9 and L4 genome size. Since our results show relatively equivalent peaks in terms of abundance, distribution length and genomic proportion, we compared raw DNA sequence peak composition (described in **Methods**) between Sf9 and L4 samples. This analysis retrieved only 11% of homologous peaks in the best case (**Supplementary Fig. 4**). In order to analyze genome-wide distribution of H3K9me2 enriched domains between samples, we proceeded to the systematic annotation of genes in both *S.frugiperda* and Sf9 genomes, but also the annotation of repeated DNA, including telomeres, rDNA loci as well as transposable elements and satellite DNA (see **Methods, Supplementary Table 1**). We produced RNA-Seq replicates for Sf9 (see **Methods**) and reanalyzed published RNA-Seq data of L4 (Orsucci et al., 2020) to distinguish a pool of inactive genes from active ones by log2(TPM) expression (**Supplementary Fig. 5** and **Supplementary Table 2**).

**Figure 1:**
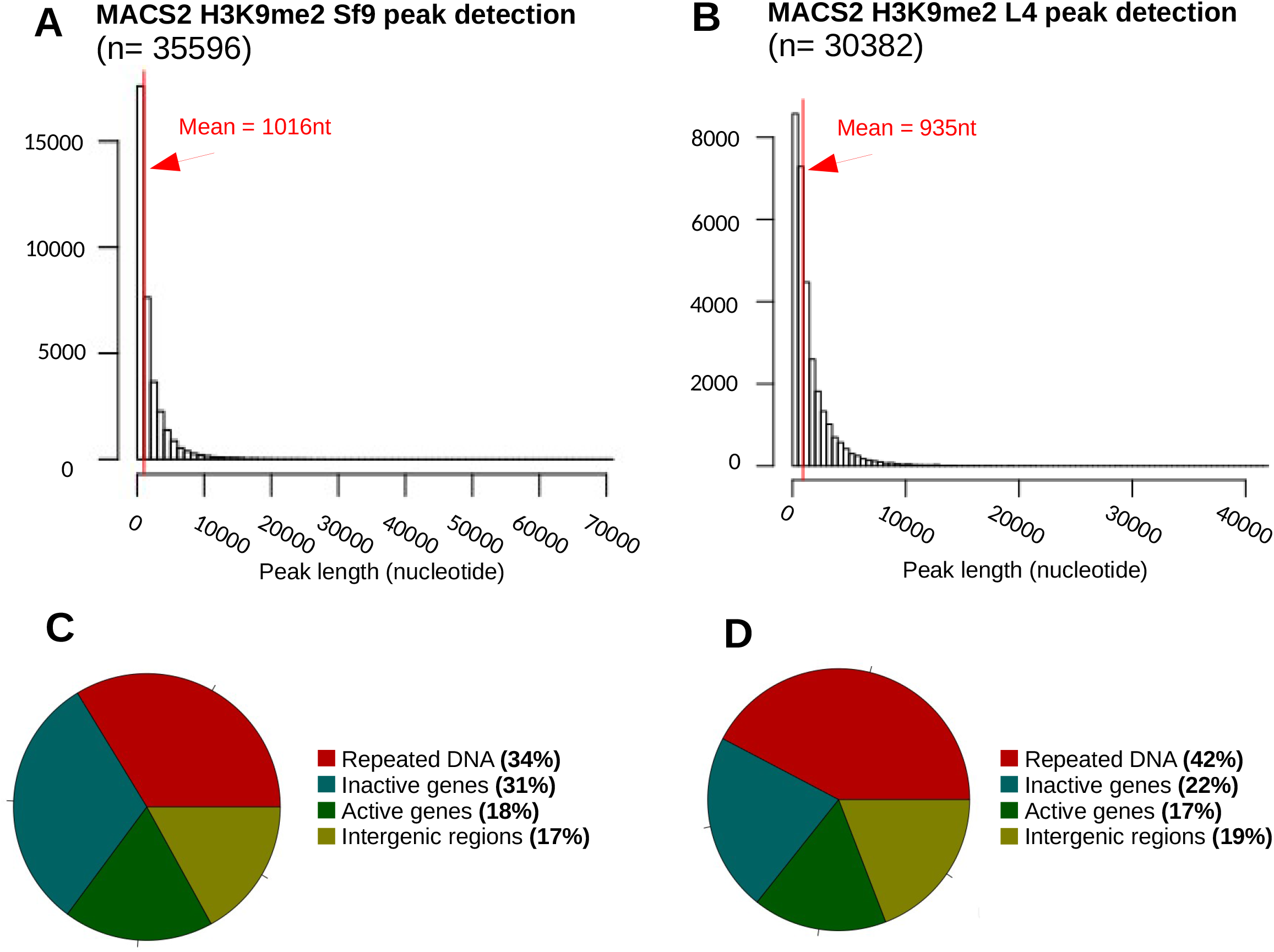
H3K9me2 genome-wide distribution in *S.frugiperda*. **A-B**: Histograms representing H3K9me2 peaks lengths detected by MACS2 in Sf9 reference (**A**) and L4 reference (**B**). **C-D**: Pie-chart showing compared abundance (%) of annotated H3K9me2 peaks (for repeated DNA, genes and intergenic regions) in Sf9 genome (**C**) and in L4 genome (**D**).

Our results show a comparable profile of H3K9me2 functional annotation over both genomes (**Fig. 1C-D**). The majority of the peaks were found associated with repeated sequences (overall of 34% and 42% in Sf9 and L4) followed by inactive genes (found respectively at 31% and 22% of H3K9me2 peaks). Surprisingly, 18% and 17% of H3K9me2 peaks were also detected within expressed genes. In addition, 17% and 19% peaks were present in intergenic regions.

While our global analysis gives clues about the general distribution of H3K9me2 in *S.frugiperda,* it failed to detect its expected association with telomeres or rDNA locus, with no enrichment found compared to broader regions such as repeated sequences and genes (**Supplementary Fig. 6**). As this result could be due to a very small fraction of telomeric and rDNA loci in the genome (~80 kb over 400 MB; **Supplementary Table 3 & 4**), we specifically analyzed the H3K9me2 distribution around the telomeric regions and rDNA locus (see Results section below) that we annotated in both reference sequences.

### 2. H3K9me2 signal in telomeres repetitions

In Noctuidae, the Lepidoptera family to which belongs the *S.frugiperda* species, chromosome pairs number is stably equal to 31 (Robinson 1971). Hence, consensual [TTAGG]n motif constituting telomeres is expected to be annotated at least 62 times in haploid genome assemblies (two tips per chromosome) (Gong et al. 2015; Vershinina, Anokhin, et Lukhtanov 2015). We searched this motif within the Sf9 and L4 genomes, and respectively detected it in 108 and 63 regions (**Supplementary Table 3**). For each presumptive telomeric region, we checked whether it was associated with any H3K9me2 peak. **Fig 2A** shows an example of homologous telomeres found in distinct Sf9 and L4 references. Homology was verified by the presence of the same upstream gene. Global analysis of these regions showed that 62 and 57 copies were enriched in H3K9me2 over the 108 and 63 annotated, representing 57±9.3% and 90±7.4% of Sf9 and L4 H3K9me2 telomere coverage (**Fig 2B**). In other biological models, ncRNA corresponding to telomeres have been found (Schoeftner et Blasco 2010). However, regardless of their length, [TTAGG]n motif sequences have almost no mapped transcripts in our samples (**Fig 2C**). Pearson correlation coefficient between RNA-Seq reads localization and telomere annotation confirms this tendency (for Sf9, R= −0.13 and for L4, R=0.002), as well as higher superior transcription state of active gene pools compared to telomeres (**Fig 2D**; Student t-test).

**Figure 2:**
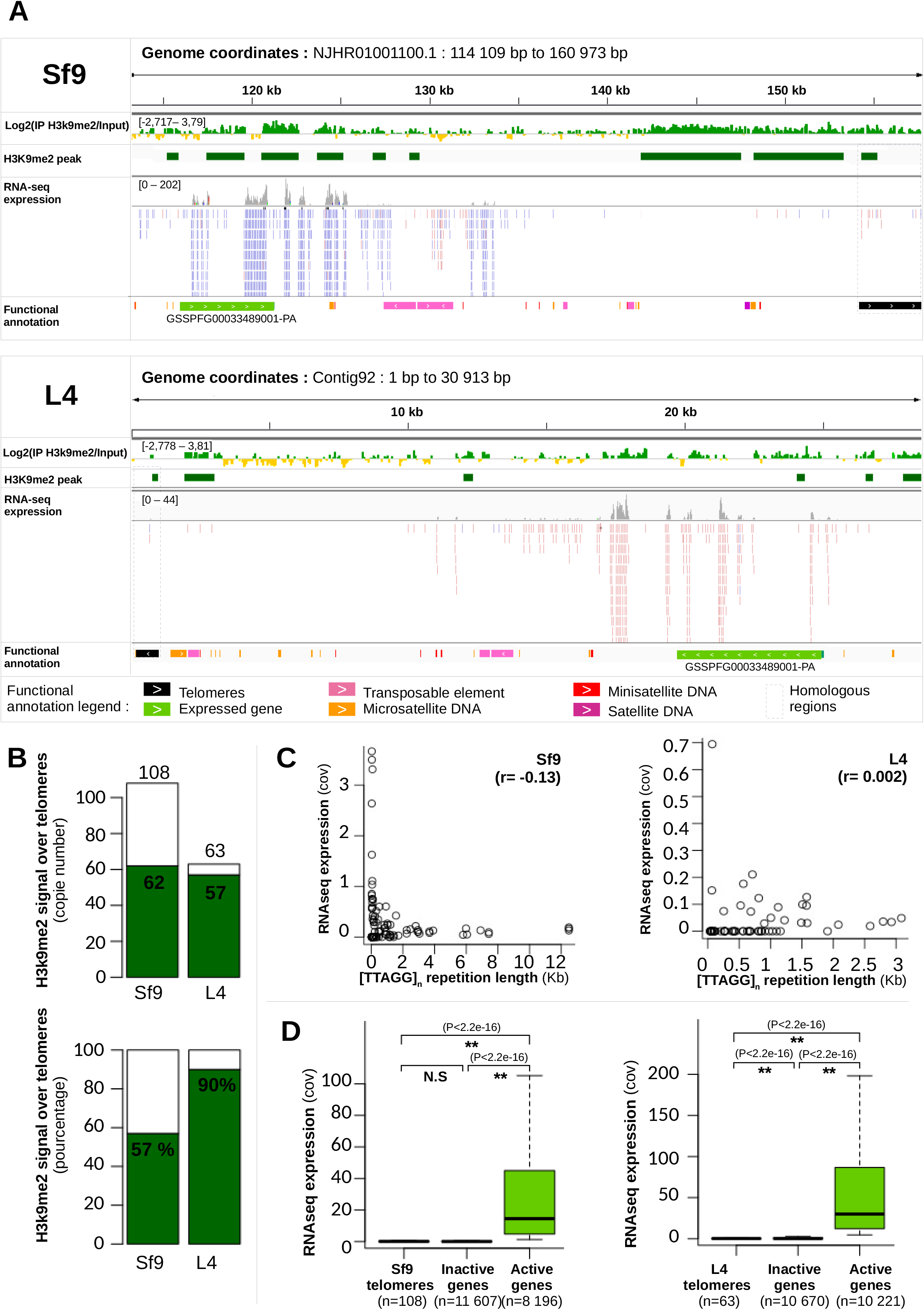
Analysis of H3K9me2 in telomeres. **A**: IGV view of homologous telomere copies found in Sf9 cells (top) and L4 (down). From the top to bottom of each view, the following tracks are displayed: 1) log2(input/H3K9me2) bigwig file (bins = 50bp), 2) H3K9me2 peaks, 3) RNA-Seq track, 4) Annotated functional element (genes, repeated DNA) **B**: H3K9me2 peak abundance in telomere annotated copies (top). Same result is shown with percentages (down). **C**: Correlation between RNA-Seq expression (in cov, y-axis) and telomere length annotated copies (in kb, x-axis). Pearson correlation for Sf9 (left) and L4(right) are both indicated. **D**: RNA-Seq expression boxplot comparison between telomeres, inactive genes and active genes are shown for Sf9 (left) and L4 (right). Statistical significance has been assessed by t-test. (*): P<0,05;(**): P< 0,001, NS: non-significant.

### 3. H3K9me2 signal in rDNA locus

Previous FISH cytology experiments conducted in Noctuidae showed the conservation of one rDNA locus located interstitially in an autosome (Nguyen et al. 2010). This polycistronic-like cluster is made of repeated 18S, 5.8S and 28S genes with additional 5S RNA being anti-sense included or extra located (Shaw et McKeown 2011). In case of low traductional activity, some rDNA copies are heterochromatinized with H3K9me2 mark (Srivastava, Srivastava, et Ahn 2016). Like we did for telomere sequences, we compared H3K9me2 enrichment and transcription at rDNA loci. We searched both reference genomes for rDNA sequences and found one major locus in both, even though relatively shorter rDNA sequences and even larger ones can be detected at various places in the genome (**Supplementary Table 4**). The homologous rDNA regions in Sf9 and L4 genomes are highly transcribed with respective mean coverage values of 78.51 X and 320.7 X, even though RNA-Seq has been performed by mRNA enrichment through ribosomal DNA depletion (**Fig. 3A**). Intriguingly, we observed in Sf9 a co-occurrence of H3K9me2 enrichment and high RNA transcription (**Fig. 3A**, upper panel). This counterintuitive result can be explained by sequence nature: since rDNA clusters are made of the same repeated sequences, short reads can align to heterochromatinized domains as well as euchromatic ones. Thus, if only one or few portions are H3K9me2 enriched in biological reality, every identical DNA sequence would be predicted as associated with the mark. The results are more coherent in L4 with RNA expression coinciding with absence of H3K9me2 peak. rDNA expression is even found overexpressed when comparing the rDNA cluster to a pool of active genes (**Fig. 3B**, Student t-test, pvalue < 2.2×10e-16).

**Figure 3:**
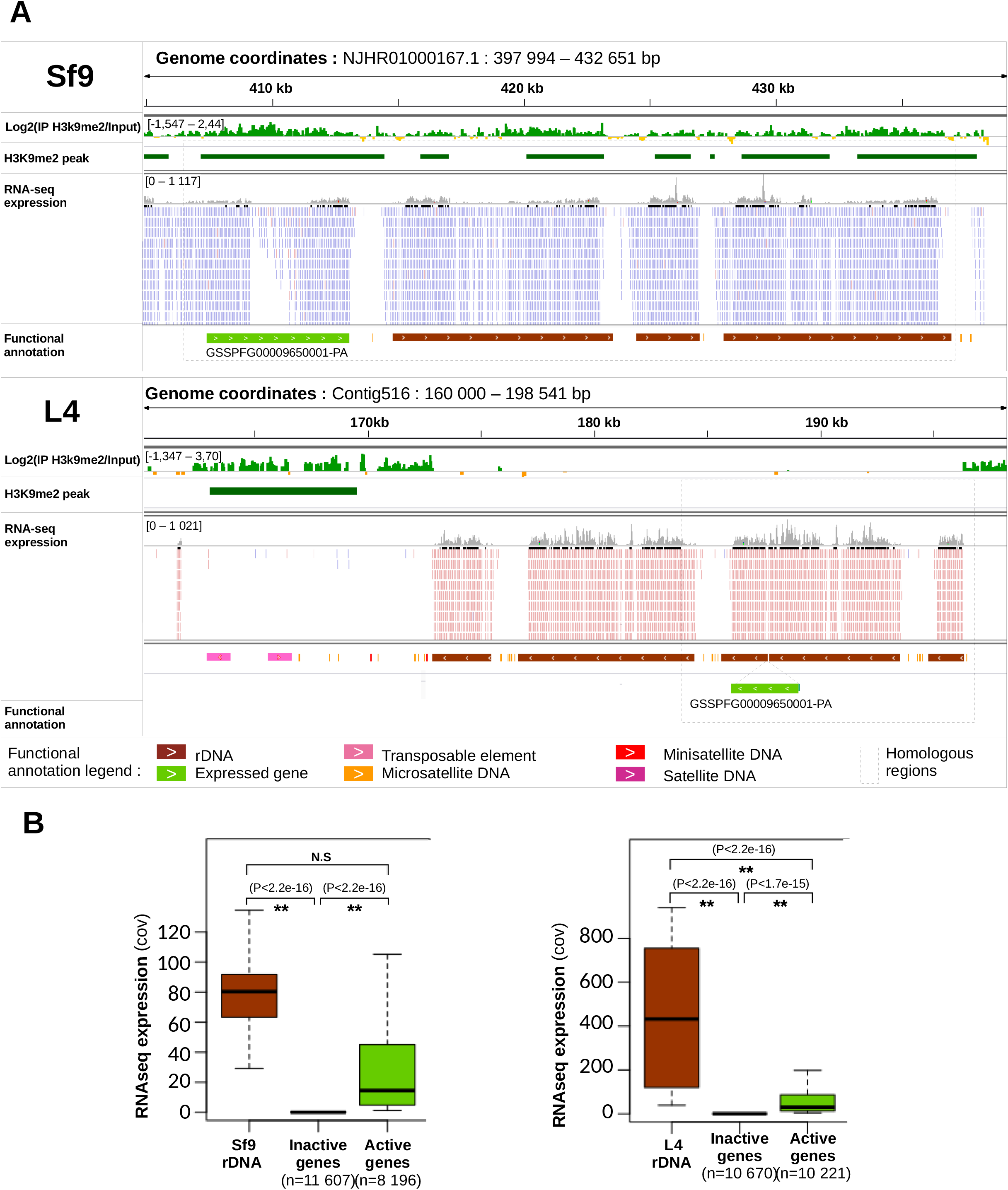
Analysis of H3K9me2 in repeated ribosomal locus. **A**: IGV view of the rDNA copies found in Sf9 cells (top) and L4 (down). Tracks from top to bottom are: 1) log2(input/H3K9me2) bigwig file (bins = 50bp), 2) H3K9me2 peaks, 3) RNA-Seq track, 4) Annotated functional element (genes, repeated DNA) **B**: RNA-Seq expression boxplot comparison between rDNA, inactive genes and active genes are shown for Sf9 (left) and L4 (right). Statistical significance has been assessed by t-test. (*): P<0,05;(**): P< 0,001, NS: non-significant.

### 4. H3K9me2 enrichment around satellite DNA regions

In many organisms, heterochromatin is associated with pericentromeric regions (Sullivan et Karpen 2004; Janssen, Colmenares, et Karpen 2018, **Supplementary Fig. 7**). These regions can be quite large, spanning several Mb of sequences (Riddle et al. 2011). We wondered whether heterochromatin in holocentric species is also associated with pericentric regions. To detect putative centromeres in *S.frugiperda* we searched and annotated satellite DNA sequences (Subirana et Messeguer 2013). Indeed, a previous molecular evolution study conducted on 282 species showed that the most abundant 150bp satellite repetitions present in the genome correspond to centromeric DNA sequences (Melters et al. 2013). This was confirmed for the majority of monocentric organisms but contested for holocentric ones (Melters et al. 2013) with only *Rynchospora pubera* plant sharing this characteristic (Marques et al. 2015). We searched the most abundant 150bp satDNA in *S.frugiperda* and analyzed its chromosomal distribution, its transcriptional status and its peripheral H3K9me2 enrichment.

Our analysis (see **Methods**) identified one 150bp satellite DNA (satDNA) consensus sequence shared between Sf9 and L4 (**Fig. 4A**). This repeated sequence is found in 1184 and 1238 copies within Sf9 and L4 genomes respectively (**Fig. 4B**). These satDNA regions do not overlap any previously annotated functional sequences (telomere, rDNA, genes etc.). Their median length is between 280 and 291 base pairs, which could correspond to 2 nucleosomes. Transcription rates were found lower or similar to annotated telomeres or the pool of inactive genes in both references (**Fig. 4C**, Student t-test). Interestingly, H3K9me2 enrichment profile around satDNA was found similar in the two references with a systematic decrease of the mark inside candidate sequences opposing a broader adjacent signal. When candidate sequences are shuffled for genomic localization, no H3K9me2 enrichment is detected in and around regions of interest (**Supplementary Fig. 8**), confirming that H3K9me2 association with satDNA is not random. In both references, highest adjacent H3K9me2 peaks are stably found around 1000bp from major 150bp satellites (**Fig. 4D**). While this result might indicate an association of heterochromatin with centromeric regions in holocentric species, additional studies would be needed to confirm that satDNA corresponds to *bona fide* centromeres in *S. frugiperda*.

**Figure 4:**
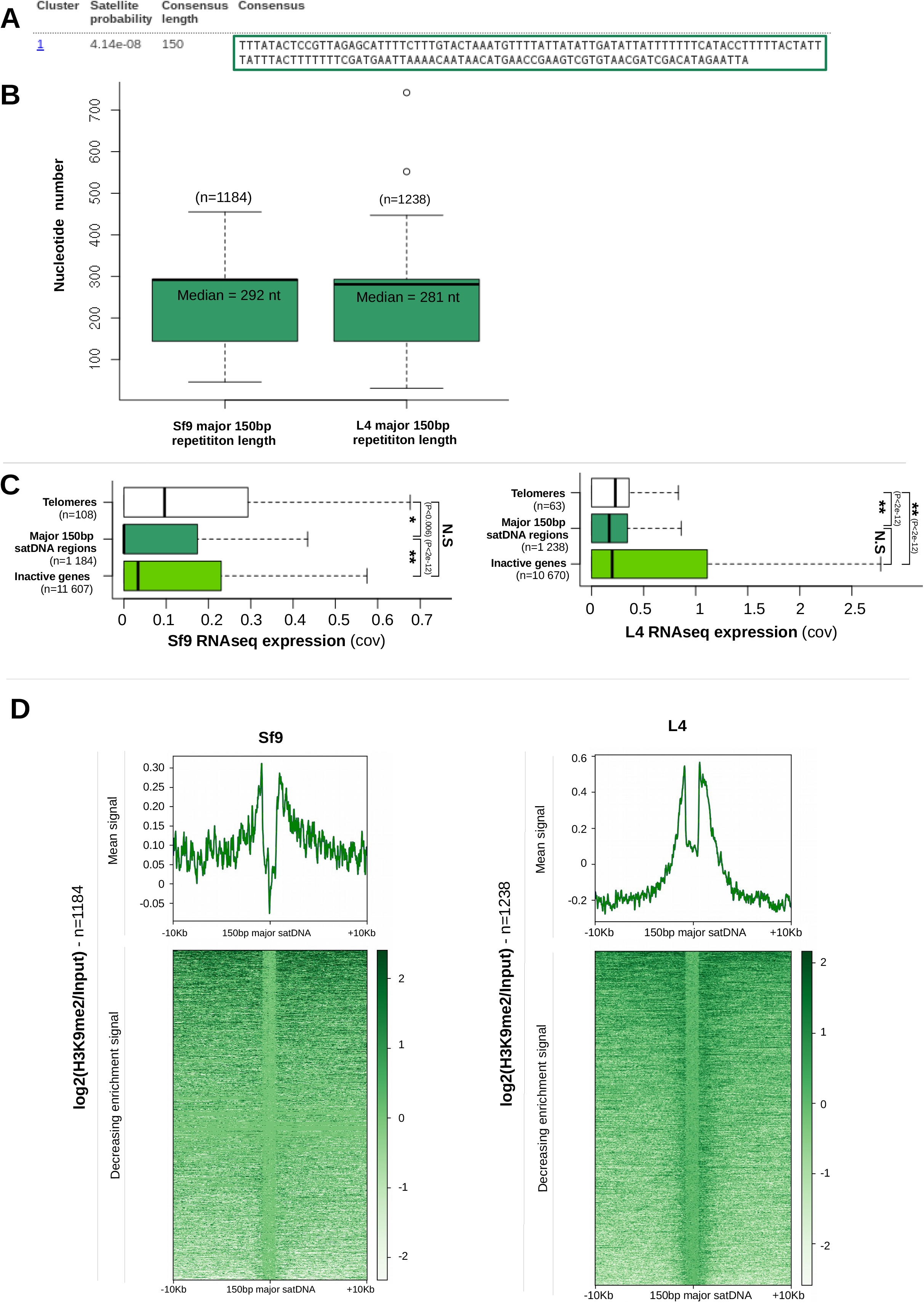
Analysis of H3K9me2 around most abundant 150bp satDNA repeat. **A**: Table showing first rank 150bp satDNA detected in Sf9 and L4 genome by RepeatExplorer. Probability, consensus length and DNA sequence composition are indicated. **B**: Boxplot of 150bp satDNA repetition regions length in Sf9 (left) and L4 (right) **C**: Boxplot of RNA-Seq expression (in cov, x-axis) between telomeres, 150bp candidate regions and inactive genes of Sf9 (left) and L4 (right). Statistical significance of expression has been assessed by t-test. (*): P<0,05;(**): P< 0,001, NS: non-significant. **D**: H3K9me2 expression of peripheral major 150bp satDNA regions in Sf9 (left) and L4 (right). For upper graphs: Mean log2(H3K9me2/Input) signal (y-axis) in 10kb regions surrounding region of interest (x-axis). For lower graphs: Decreasing log2(H3K9me2/Input) expression (y-axis) of 150bp satDNA regions and 10kb around (x-axis)

### 5. H3K9me2 enrichment in repeat elements families

Between 34% to 42 % of H3K9me2 peaks are associated with repeat sequences (**Fig. 1C,D**). We annotated in both reference genomes the different categories of repeat sequences, whether they correspond to tandem repeats, such as micro- and mini- satellite sequences, or transposable elements (**Fig. 5A**). We found around a hundred thousand micro- and minisatellite sequences in both Sf9 and L4 genome assemblies, representing 3 674 661 2 875 505bp and 1 706 018 1 149 269 bp each, or about 0.7% and 0.3% of the genome. We found less satellite sequence repeats (1719 and 3067 sequences) representing 0.5% and 0.2% of the genome but a large amount of putative transposable elements (46 625 and 55928 sequences representing 6.7% and 9.8% of the genome).

**Figure 5:**
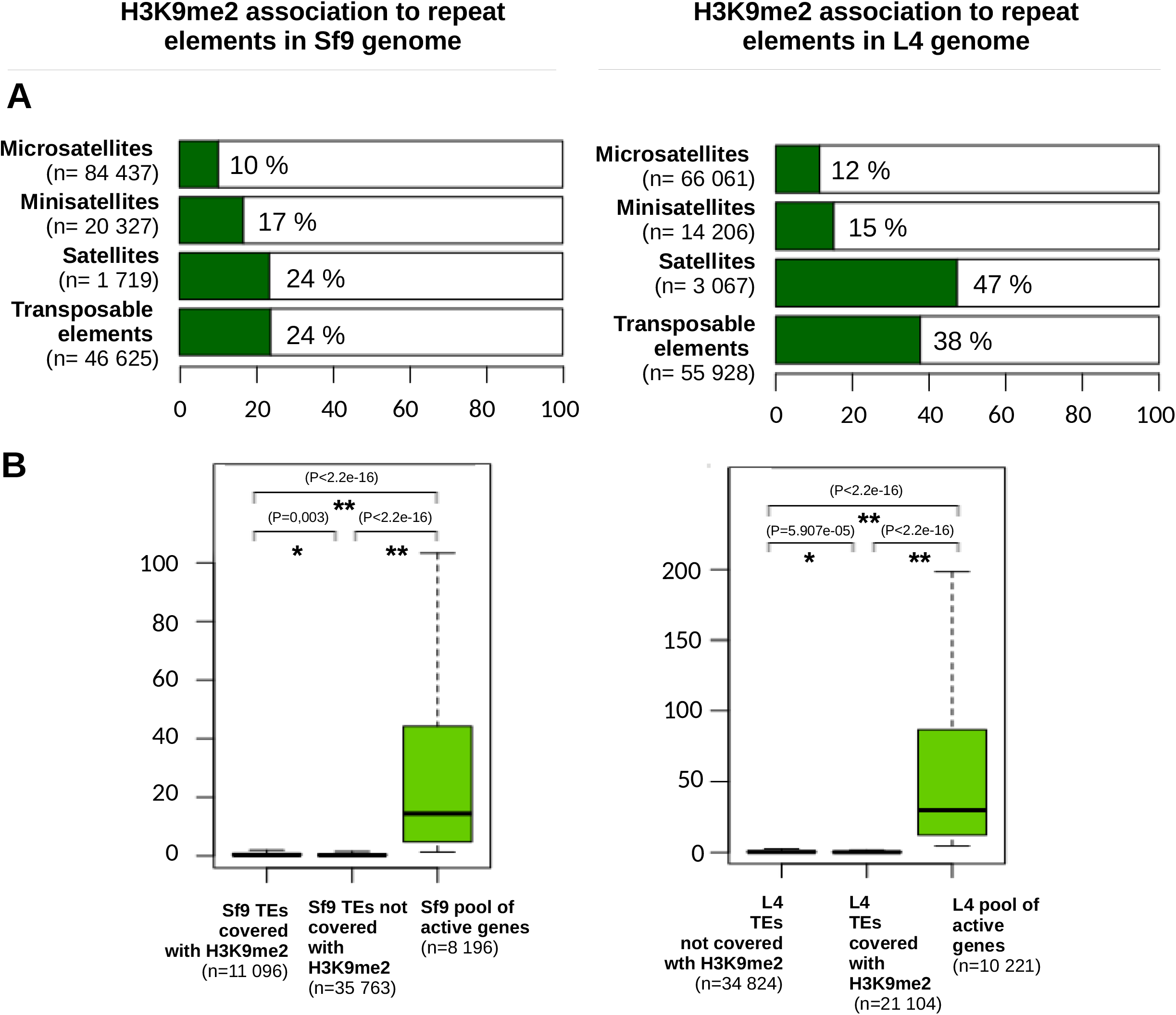
Analysis of H3K9me2 enrichment with repeat element families. **A**: Plots showing abundance (in %, x-axis) of H3K9me2 in annotated tandem repeats and transposable elements of Sf9 (left) and L4 (right) genomes **B**: Boxplot comparing transposable elements RNA-Seq expression between those covered with H3K9me2, others without epigenetic mark association and pool of active genes

We then calculated the proportion of each category associated with H3K9me2 (**Fig. 5A**). A higher proportion of satellite sequences and transposable elements is found associated with H3K9me2 compared to micro- and mini-satellite, which agrees with our expectation of a role of heterochromatin associated with repeat-rich regions of the genome.

The higher prevalence of repeat DNA and transposable elements in c-Het compartments in model organisms is often explained by the repressive nature of heterochromatin. Indeed, because of their potential deleterious effects on the genome when they are mobilized, transposable elements are often transcriptionally repressed by small RNA targeted H3K9me (Klenov et al. 2007; Sienski, Dönertas, et Brennecke 2012; Le Thomas et al. 2013). In order to determine if a similar role of H3K9me2 exists in *S. frugiperda* we analyzed the RNA-Seq data from L4 tissues and Sf9 cells to classify transcribed and non-transcribed transposable elements and observe the association of each category with H3K9me2 signal (**Fig. 5B**). No enrichment was observed, with H3K9me2 being equally found between active and inactive transposable elements.

### 6. H3K9me2 signal enrichment in genes

Due to the classical repressive nature of heterochromatin, we expected H3K9me2 to be associated with mainly inactive genes. However, our ChIP-Seq analysis detected a comparable H3K9me2 enrichment in expressed genes compared to inactive ones in both references (**Fig. 1B**).

We analyzed if some gene regions like promoters, UTRs or gene bodies were more enriched in H3K9me2 than others (**Fig. 6A**). We found inactive promoters to be statistically more associated with H3K9me2 than active promoters in the two references (Sf9: χ^²^ = 317.37, df = 1, p-value<2.2e-16; L4: χ^²^ = 667.97, df = 1, p-value<2.2e-16). However, 16% Sf9 and 26% L4 active promoters were also associated with H3K9me2 marks, which may be due to the cell heterogeneity in both samples or to difficulties in promoter prediction in our model. H3K9me2 is also detected within the gene body and UTRs regardless of the gene transcriptional status in equivalent proportions between Sf9 and L4 cells (**Fig. 6A**). Interestingly, when we took a further look into differently H3K9me2 enriched for gene body regions, we observed a more intense signal in exons compared to introns.

**Figure 6:**
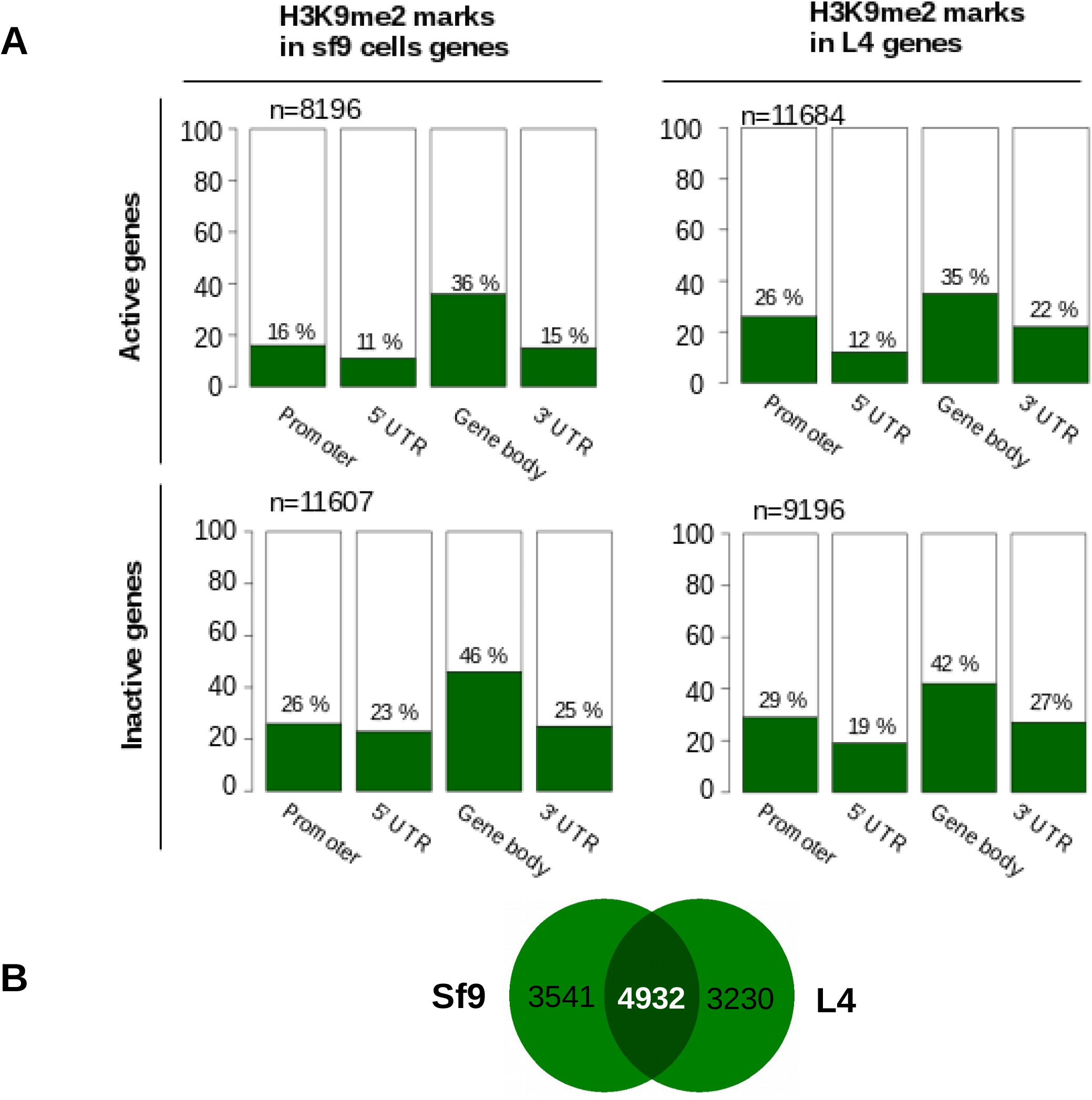
Analysis of gene regions covered with H3K9me2 mark. **A**: Plots showing abundance (in %, y-axis) of H3K9me2 present in promoters, UTRs and CDS (x-axis) of active genes (upper part) vs. inactive genes (lower part) from Sf9 (left) and L4 (left) cell models. **B**: Venn diagram showing unique genes respectively covered in Sf9 and L4 (left and right part) and common ones (middle part)

In Sf9 cells, 42.8% of annotated genes are associated with H3K9me2 and 39.1% in L4 (**Fig. 6B**). For those genes, H3K9me2 signal is distributed within exons rather than introns (**Fig. 7**). Two-thirds of the H3K9me2 covered genes regardless of their transcription state are shared between the two references (**Fig. 6B**). For these common genes, we performed a Gene Ontology analysis. We observed an enrichment in functions associated with transposable elements regulation (**Supplementary Fig. 9**, GO: endonuclease activity, nuclease activity) and nucleic acid homeostasis (**Supplementary Fig. 9**, GO: DNA metabolic process, catalytic activity acting on DNA etc.). We also noticed the presence of genes involved in heterochromatin maintenance and formation, suggesting possible molecular feedback. It includes a H3K9me methyltransferase enzyme (Suvar3/9: GSSPFG00004579001-PA, G9a GSSPFG00019340001.2-PA), DNA methylase enzyme (DNMT1 (GSSPFG00025486001.2-PA)), centromere formation proteins (search from Cortes-Silva et al. 2020; inner kinetochore: CENP-P (GSSPFG00001205001-PA)), ATP synthase subunit (GSSPFG00010096001-PA; Collins et al. 2018), outer kinetochore:Nuf2 (GSSPFG00010779001-PA), BLAST2Go annotated centromeric proteins (GSSPFG00020169001-PA, GSSPFG00005386001.4-PA)) and HP1 family proteins (HP1c (GSSPFG00011657001.2-PA),HP1e (GSSPFG00007777001.2-PA)). Paradoxically, lots of spliceosome genes involved in active transcription and mRNA formation are also found.

**Figure 7:**
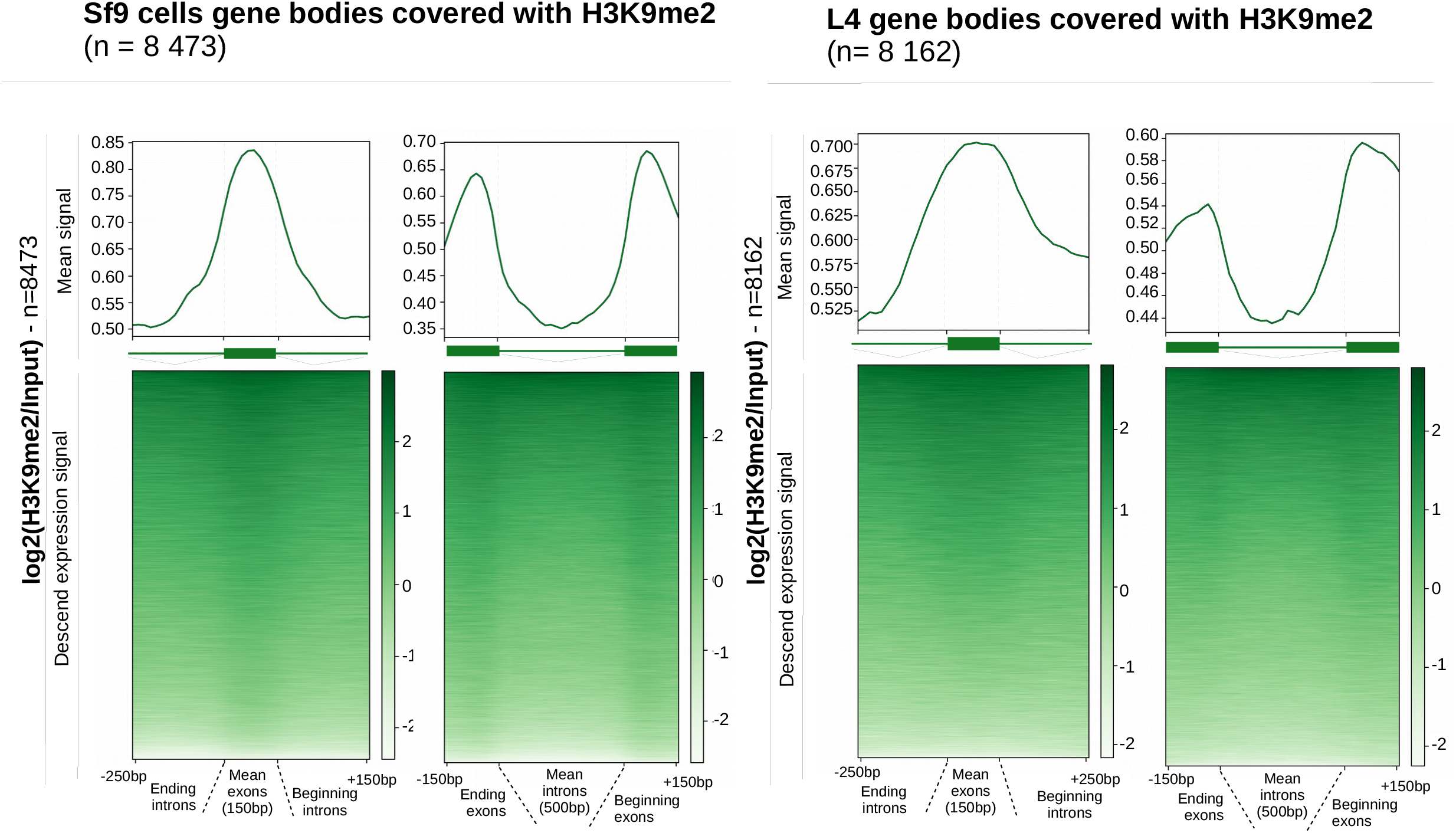
Analysis of H3K9me2 signal covering exons vs introns gene bodies. Left: Sf9; Right: L4. For upper graphs: Mean log2(H3K9me2/Input) signal (y-axis) in all exons (left) and introns (right, x-axis). For lower graphs: Ascend log2(H3K9me2/Input) expression (y-axis) in all exons (left) and introns (right, x-axis).

## Discussion

We report the first H3K9me2 genome-wide ChIP-Seq analysis conducted in *S. frugiperda* a Lepidopteran species. Our results are globally consistent with previous studies on H3K9me2 (Ho et al. 2014) albeit with a higher percentage of genome covered (13.8±0,02% and 12,6±0,03% in Sf9 and L4) than in other monocentric and holocentric organisms. We detected H3K9me2 at expected chromosomal compartments such as telomeres and rDNA locus but no major centromeric locus. Instead, we observed a scattered distribution of H3K9me2 along the chromosomes colocalizing with genes and repeat elements, independent of their transcriptional status.

### Localization of H3K9me2 at expected heterochromatin compartments

In many monocentric organisms, c-Het is found in large regions surrounding the centromere. Since we found H3K9me2 to be still associated with heterochromatin domains such as telomeres or rDNA locus in *S.frugiperda* (**Fig. 2,3**), we hypothesized that c-Het could also identify putative centromeric regions in this holocentric specie. As mentioned in the introduction, H3K9me2 surrounding the major 150bp satDNA repetition is a conserved pattern for pericentric regions in monocentric species. To retrieve putative centromeric regions in the genome of *S.frugiperda* we annotated the most abundant 150bp satDNA and found it present in more than a thousand scattered copies in both genomes (**Fig. 4**). Since we also observed an enrichment of H3K9me2 within 1kb of satDNA, we hypothesize they could represent holocentromeres. A recent publication on Lepidopteran cell lines (*Bombyx mori* and *Trichoplusia ni*) describes centromeres marked by the centromeric protein CenP-T to be unusually associated with the facultative heterochromatin mark H3K27me3 (Cortes-Silva et al. 2020; Senaratne et al. 2021). However, the distribution of H3K9me2 was not assessed in their study and we don’t know whether it colocalizes with H3K27me3 or is excluded. It is possible that satDNA sequences associated with H3K9me2 represent in fact vestigial centromeres that stopped being used by the cell after H3K27me3 replacement. To determine if such genomic regions correspond to functional holocentromeres, its association with kinetochore proteins would have to be demonstrated.

### Repeated sequences and H3K9me2 association

Among the conserved features of H3K9me2 distribution, we observed this mark to be mainly associated with repeated DNA sequences in both cellular references. This was evidenced by the high abundance of multimapper reads in both cellular references (**Table 1**) but also by the significant ChIP enrichment on the different categories of annotated repeated DNA (**Fig. 1C**). While this association was expected from previous studies conducted in Lepidopteran (Stanojcic et al. 2011; Borsatti et Mandrioli 2013), we present here a more exhaustive association with the different categories of repeats. We did not observe a strong complete association of the mark with repeat elements with only 24% and 38% of transposable elements intersecting H3K9me2 peaks (**Fig. 5A**). In addition we did not observe a clear correlation between transposable elements repression and H3K9me2 as we were expecting (Klenov et al. 2007, **Fig. 5B**). It is possible that other epigenetic marks might be involved to specifically repress transposable elements Future genome-wide H3K27me3 or H3K9me1/3 ChIP-Seq data should be compared with H3K9me2 to determine if, in Lepidoptera, transposable elements control is insured by similar heterochromatin mechanisms as in other model insects.

### H3K9me2 association with genes bodies

H3K9me2 in *S.frugiperda* is associated with some gene bodies regardless of their active or inactive transcriptional status (**Fig. 1B**). This histone mark has been described to cover coding sequences in two different situations.

The first situation corresponds to hundreds of genes that are present in c-Het compartments. Contrary to silenced euchromatic genes at heterochromatin proximity -an effect called position effect variegation (PEV, Wallrath et Elgin 1995)- these heterochromatic genes do not function when displaced in euchromatin (Yasuhara et Wakimoto 2008; Dimitri et al. 2009; Saha et al. 2020). Their expression is compatible with H3K9me2 covering. Principally described in drosophila, some of these genes are essential for development and are mainly concentrated in pericentromeric regions. Given their localization, they are thought to be more prone to transposable elements insertions and show lots of intronic transposons. In *S.frugiperda* we identified more than 4900 genes associated with the H3K9me2 mark. Their function reflects constitutive roles such as general transcription factors, ribosomal genes or mitochondrial genes, which fits with heterochromatic genes hypothesis.

The second situation described in the literature of H3K9me2 association with gene bodies is in alternative splicing. Presence of exonic H3K9me2 across genes is thought to slow down polymerase in order to favor alternative instead of constitutive mRNA transcription (Saint-André et al. 2011). Consistent with this, our analysis of H3K9me2 signal in *S.frugiperda* shows a higher enrichment in exons compared to introns (**Fig. 7**). In order to confirm a splicing role for this mark in *S.frugiperda* it would be necessary to annotate exons nature (internal vs. external, constitutive vs. alternative, etc.) and check if alternative mRNA transcripts are produced given presence or absence of exonic H3K9me2 signal. If previous studies focused on other heterochromatic components such as DNA methylation or HP1c proteins to regulate transcription and alternative splicing in insects (Bonasio et al. 2012; Li-Byarlay et al. 2013), no studies linking these epigenetic factors with H3K9me2 has been addressed in Lepidopteran. This fact is important since HP1c is usually described on both exons and introns. A recent work revealed intronic HP1c signal to be involved in alternative splicing through binding with abundant “CACACA” intronic repeated motif sequences (Rachez et al. 2019). Our search for a similar motif in *S.frugiperda* using two dedicated softwares failed to give the same results. But a systematic association of HP1c and H3K9me2 cannot be assessed since HP1c can bind to other modified marks and RNAs. In addition, the correspondence of classical HP1 proteins with homologs in Lepidoptera is not clear. In *Bombyx mori* two HP1 homologs have been described: BmHP1a and BmHP1b (Mitsunobu et al. 2012). We also retrieved those two homologs in *S.frugiperda* genomes even though the phylogeny needs to be more resolved since they are not on the same branch as the classical Drosophila HP1a (**Supplementary Fig. 10**).

## Conclusion Article summary

We produced the first genome-wide analysis of holocentric Lepidoptera *S. frugiperda* heterochromatin distribution by analyzing H3K9me2 ChIP-Seq data in two cell models. In contrast to studies suggesting unusual behavior of modified histones, our results supported a conserved pattern with invariant classic c-Het domains such as (sub)telomeres, rDNA locus and even peripheral major 150bp satDNA that could be associated with centromeric functions. However, we also advocate for a more pleiotropic function of H3K9me2 since it is abundantly present in transposable elements as well as gene bodies regardless of their expression status. In order to deepen these results and get a more detailed picture of heterochromatin localization in holocentric Lepidoptera, further work characterizing other associated histone marks, DNA methylation and HP1 proteins genome-wide distribution in Lepidoptera will be required.

## Acknowledgements

This work was supported by a grant from the French National Research Agency (ANR-12-BSV7–0004-01; http://www.agence-nationale-recherche.fr/) for EA and Institut Universitaire de France for NN. The funding bodies had no role in the design of the study and collection, analysis, and interpretation of data and in writing the manuscript.

## Authors Contributions

SN, EA and NN designed the project and the experiments, SG, SN and RNS performed the Western-Blot, Immuno-fluorescence and ChIP experiments, DS produced the Illumina libraries and sequencing. SN produced the bioinformatic and statistical analysis with the help of KWN. SN and NN wrote the manuscript with the input of all co-authors.

## Supplementary Information

**Supplementary Fig. 1:** Immunofluorescence pictures showing H3K9me2 (red) distribution inside the nucleus at different stages of Sf9 cell cycle. Blue color is for DAPI staining and cells have been observed with a DIC microscope.

**Supplementary Fig. 2:** Western blot analysis with anti H3K9me2 antibodies of Sf9 cells (left) or L4 larvae (right) chromatin extracts. The 17KDa band corresponds to H3K9me2. Compared to Western-Blot performed with protein extracts on the same material showing one single band, extra bands here are most likely produced by cross-linking procedures.

**Supplementary Fig. 3:** Correlogram of bigwig files produced from each Input-seq and ChIP-seq samples in Sf9 and L4. Scores are Pearson correlation with 1 being most correlated and 0 least.

**Supplementary Fig. 4:** D-genies plot comparison between H3K9me2 peaks sequences between Sf9 and L4

Upper part: D-genies graph comparison between Sf9 peaks against L4 peaks (left) and L4 peaks against Sf9 ones (right). Best matches shown only.

Lower part: Summary of % sequences identity

**Supplementary Fig. 5** Histogram of log2(tpm) count associated to genes annotated in each model reference genome. Left: Sf9; Right: L4. Sf9 cells show a typically bimodal distribution representing the sets of expressed and not expressed genes. L4 larvae contain a more heterogenous cell population and have a larger set of expressed genes. Tpm: transcripts per million base pairs.

**Supplementary Table 1:** Number of functional elements annotated in this study from both Sf9 cell line and L4 larvae reference genomes.

**Supplementary Table 2**: Sf9 cells and L4 larvae RNA-seq replicates and mapping statistics. Reads abundance and Bowtie2 alignment details are described for each sample.

**Supplementary Fig. 6:** Pie chart representing H3K9me2 peaks distribution with respect the annotated functional elements for each genome reference. If a large peak encompasses different features such as an intron or an exon, we counted both elements in the distribution.

**Supplementary Fig. 7:** Schematic representation of centromeric chromatin organization in monocentric model organisms.

**Supplementary Fig. 8:** H3K9me2 enrichment signal around shuffled regions produced from major 150bp satDNA domains in Sf9 (left) and L4 (right).

For upper graphs: Mean log2(H3K9me2/Input) signal (y-axis) in 10kb (left) and 1kb (right) regions surrounding region of interest (x-axis).

For lower graphs: Ascend log2(H3K9me2/Input) expression (y-axis) of 150bp satDNA regions and in 10kb (left) and 1kb (right) periphery (x-axis)

**Supplementary Fig. 9:** Gene ontology enrichment analysis of common H3K9me2 associated genes between Sf9 and L4 models (Fisher exact test method). Only 15 most abundant category are shown (x-axis). Sequence abundance (in %, y-axis) of common vs. All annotated genes is represented.

**Supplementary Fig. 10:** Phylogenic tree of *Bombyx mori Drosophila melanogaster* and *Spodoptera frugiperda* HP1 family proteins containing a chromodomain and a chromo shadow domain. The chromodomain protein Polycomb (PC) is represented as an outgroup. *D. melanogaster* HP1 proteins (HP1a is Suvar205; HP1b, HP1c, HP1e and Pc) were retrieved from Flybase (https://flybase.org/). *S. frugiperda* proteins from L4 genome annotations start with the ‘GSSPFG’ nomenclature and have been retrieved after blastp with Drosophila HP1 proteins. *B. mori* proteins have been retrieved from ncbi (https://www.ncbi.nlm.nih.gov/protein/) after blastp with drosophila HP1 proteins.

**Supplementary Table 3**: Annotation and coordinates of putative telomeric regions in *S. frugiperda* genome based on [TTAGG]n repetitions.

**Supplementary Table 4**: Annotation and coordinates of presumptive rDNA locus in S. frugiperda genome.

