## Supplementary Material for "H3K9me2 genome-wide distribution in the holocentric insect *Spodoptera frugiperda* (Lepidoptera: Noctuidae)"

### Supplementary Fig. S1

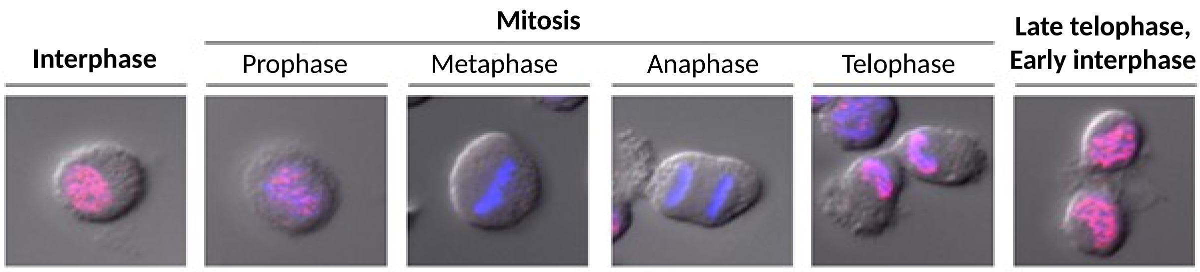

### Supplementary Fig. S2

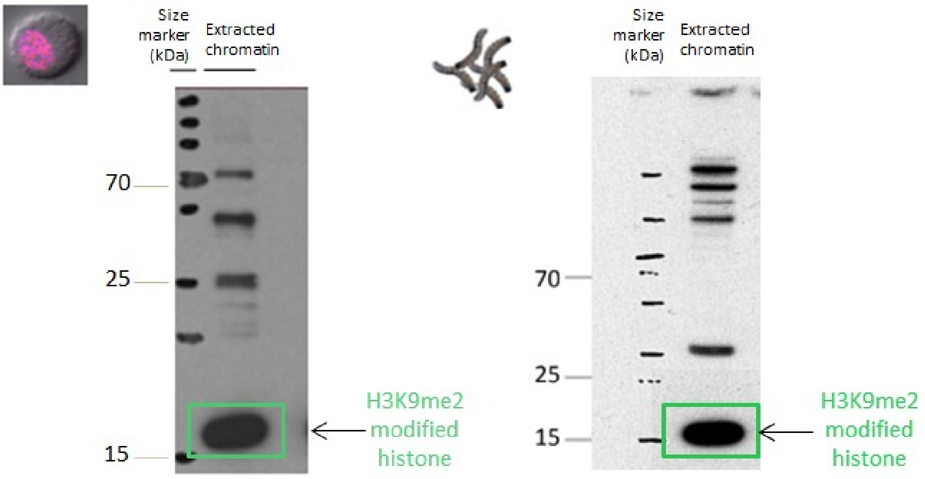

### Supplementary Fig. S3

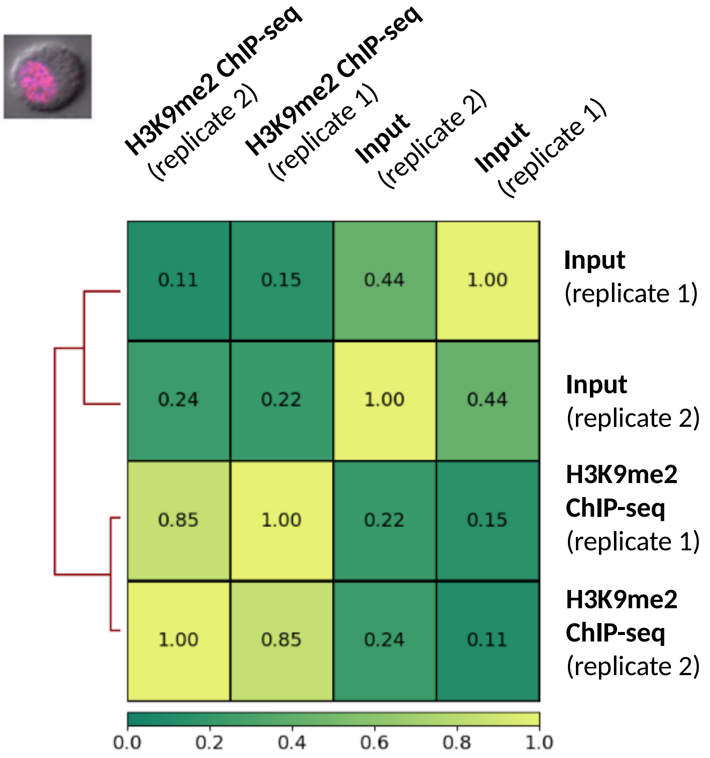


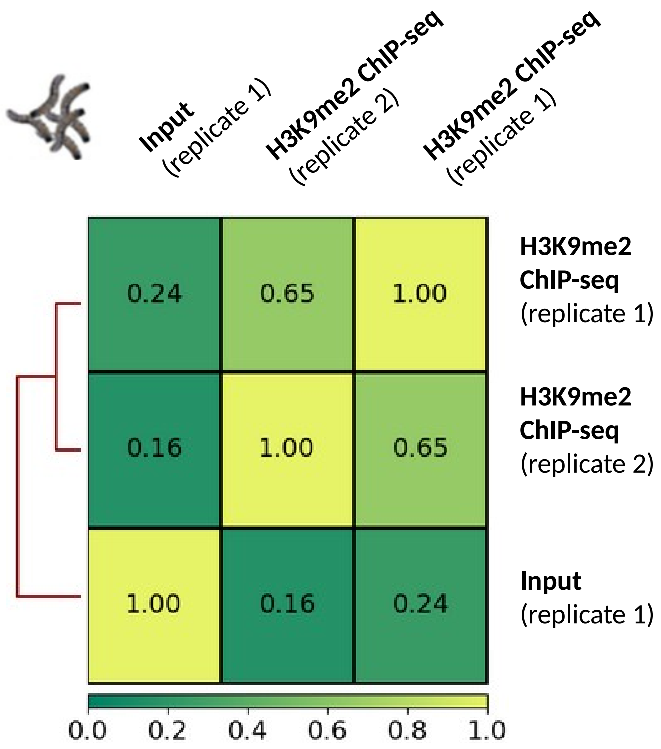

### Supplementary Fig. S4

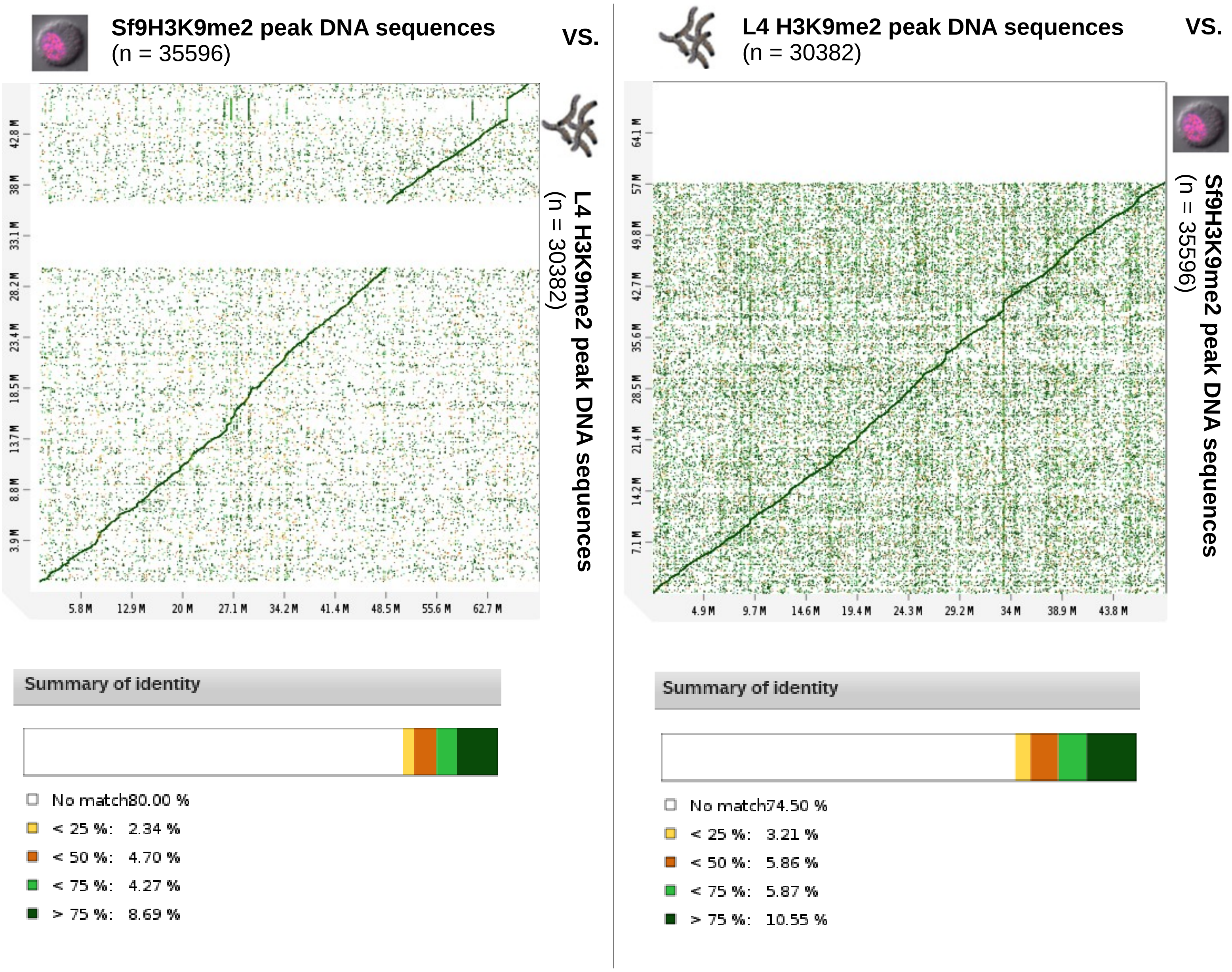

### Supplementary Figure S5

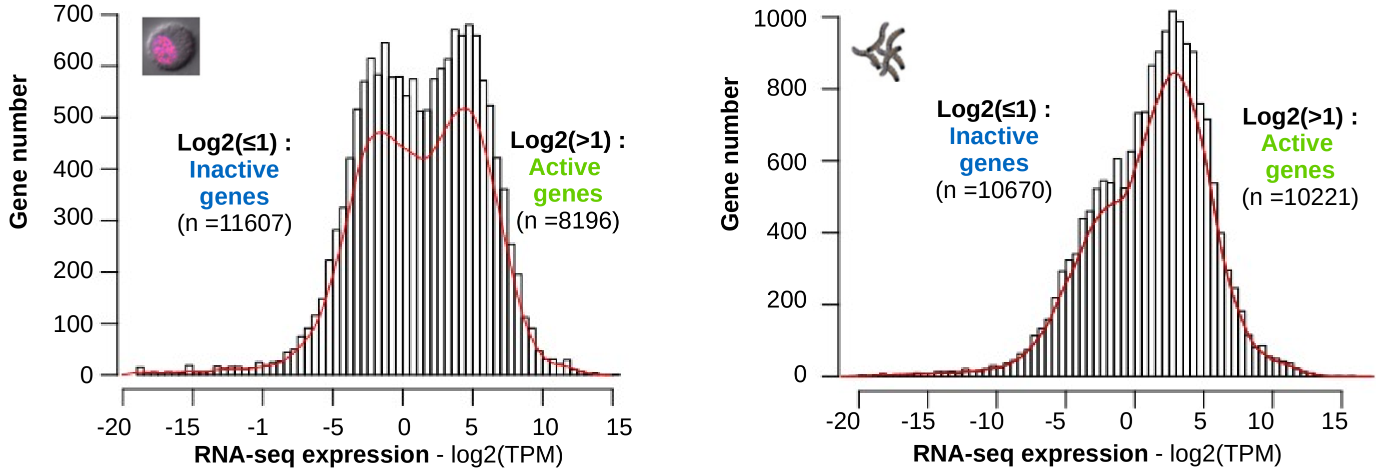

### Supplementary Table S1

| Annotations | Sf9 genome | L4 genome |
|---|---|---|
| **Scaffold number** | 2396 | 1000 |
| **Telomeres copies** | 108 | 63 |
| **rDNA repetitions** | 165 | 10 |
| **Transposable elements** | 46609 | 55928 |
| **Satellites DNA** | 1718 | 3067 |
| **Minisatellites DNA** | 20323 | 14206 |
| **Microsatellites** | 87437 | 55928 |
| **Inactive genes** | 11607 | 10670 |
| **Active genes** | 8196 | 10221 |
| **Major 150bp satDNA** | 1184 | 1238 |

### Supplementary Table 2

| Cellular model | Experimental condition | Reads number | Unmapped reads | Reads that mapped 1x | Reads that mapped >1x | Alignement rate |
|---|---|---|---|---|---|---|
| Sf9 cells 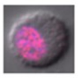 | RNA-seq (1st replicate) | 36 812 923 | 11.32% | 49.62% | 39.07% | **88.68%** |
| | RNA-seq (2nd replicate) | 36 011 772 | 11.63% | 48.36% | 40.01% | **88.37%** |
| L4 larvae 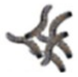 | RNA-seq (1st replicate) | 53 957 483 | 12.88% | 74.95% | 12.17% | **87.12%** |
| | RNA-seq (2nd replicate) | 34 744 137 | 12.14% | 75.64% | 12.22% | **87.86%** |

### Supplementary Figure 6

#### Sf9 cells H3K9me2 genomic distribution

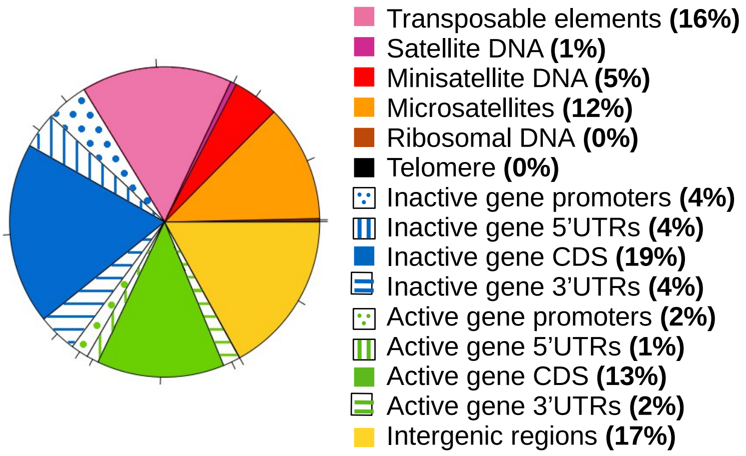


#### L4 H3K9me2 genomic distribution

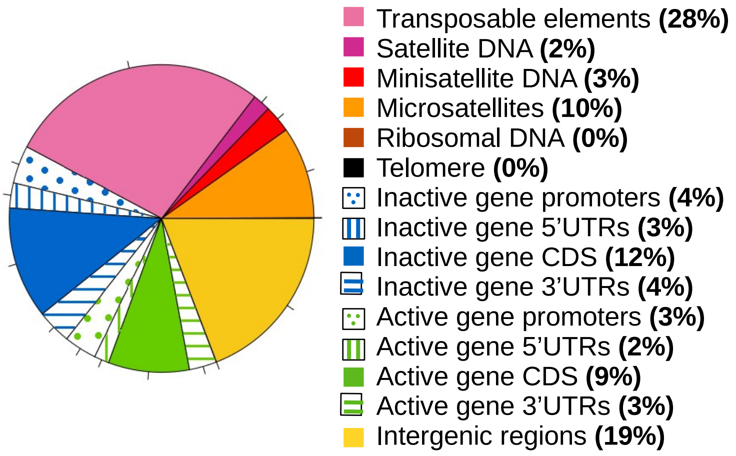

### Supplementary Figure 7

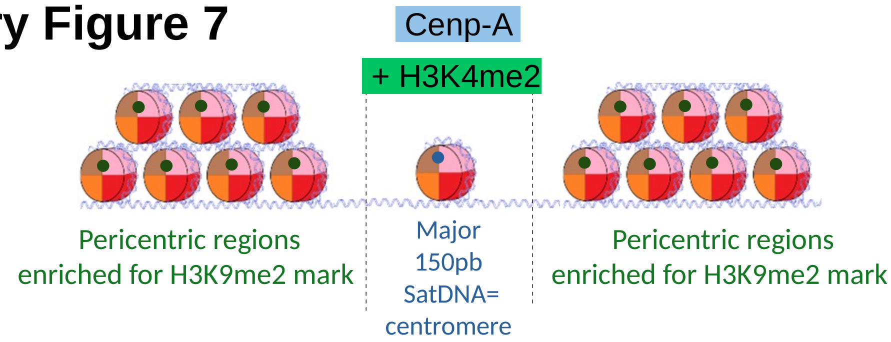

### Supplementary Figure 8

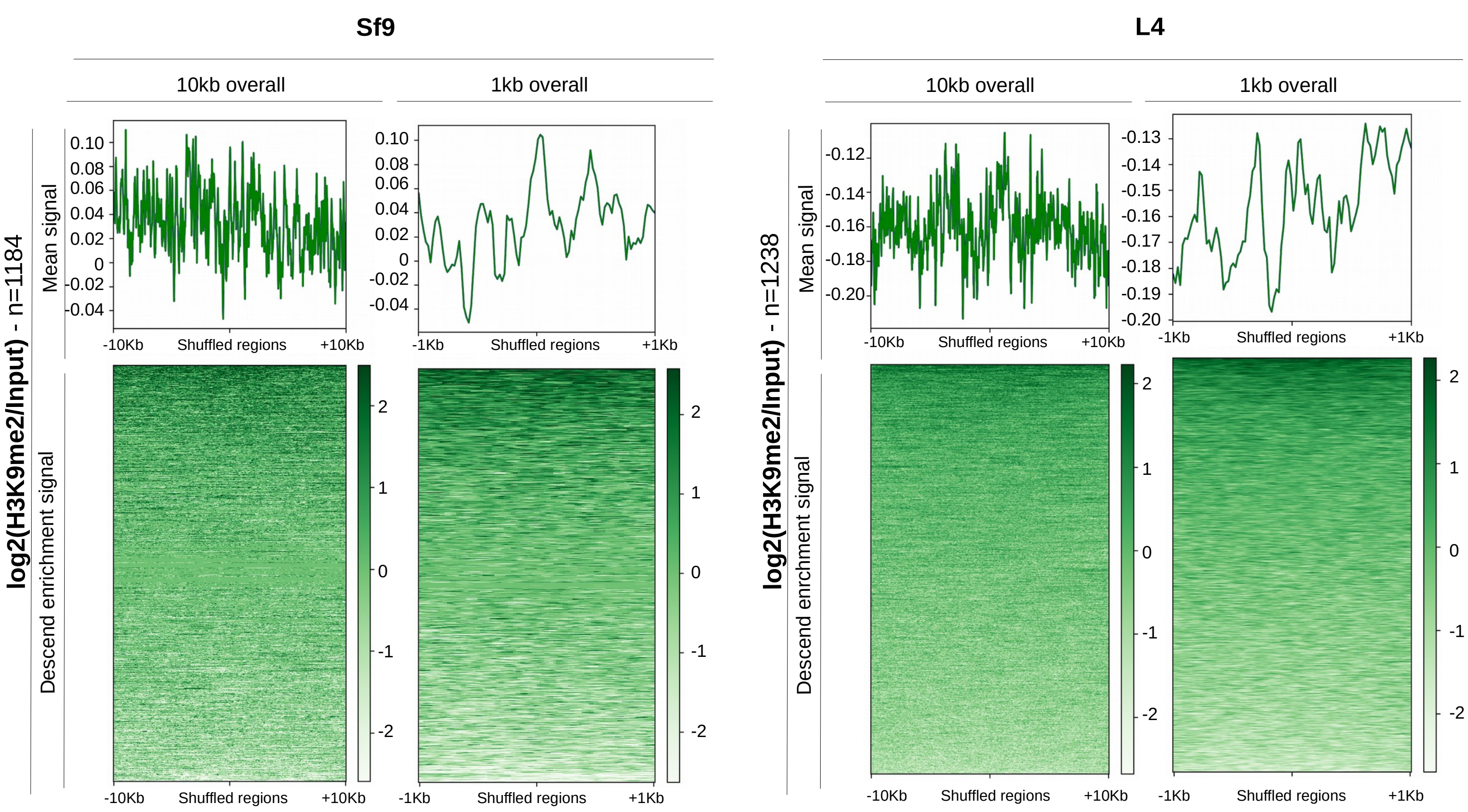

### Supplementary Figure 9

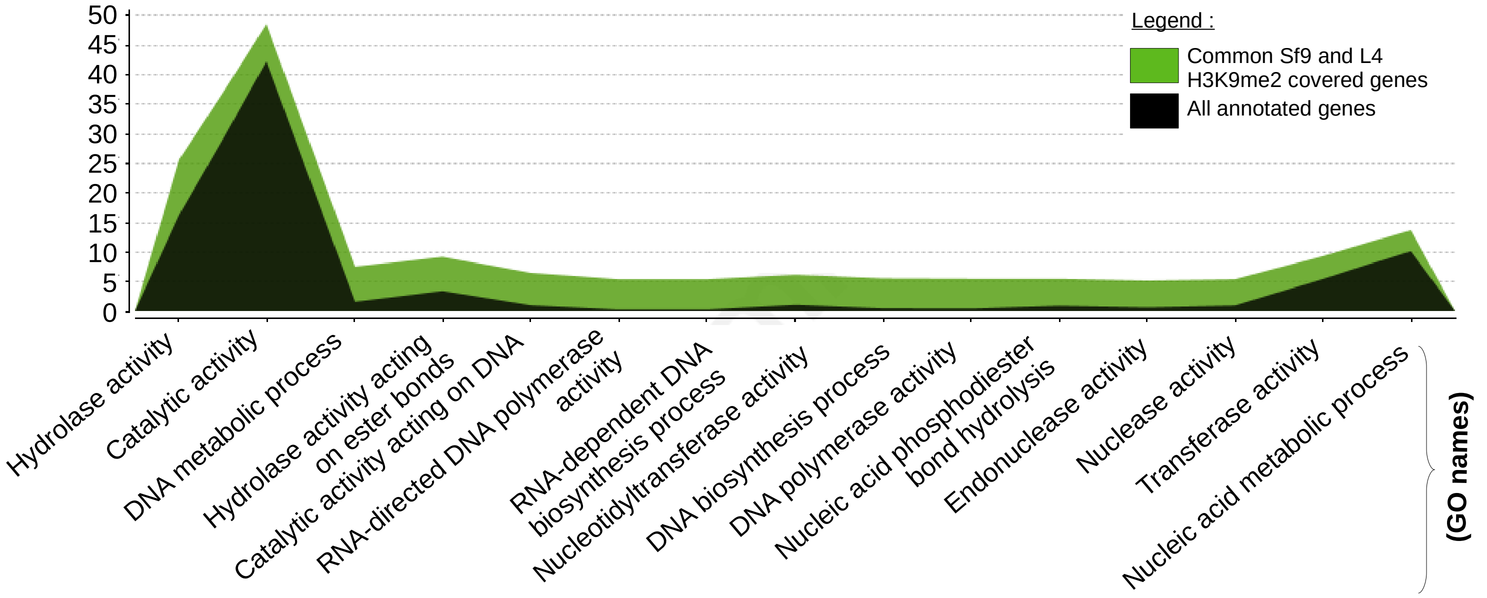

### Supplementary Figure 10

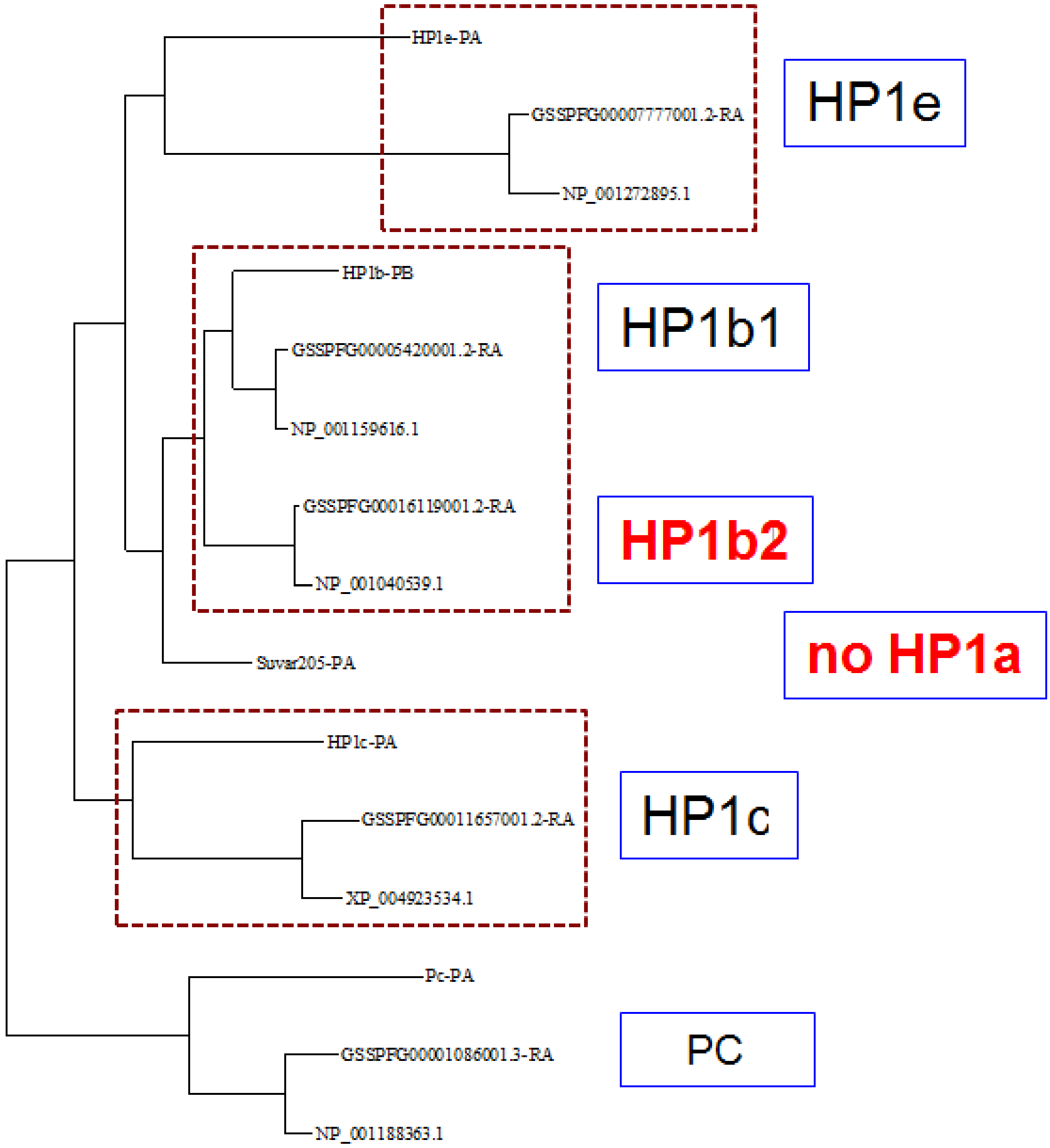
